# Chromosome-level genome assembly of a benthic associated Syngnathiformes species: the common dragonet, *Callionymus lyra*

**DOI:** 10.1101/2020.09.08.287078

**Authors:** Sven Winter, Stefan Prost, Jordi de Raad, Raphael T. F. Coimbra, Magnus Wolf, Marcel Nebenführ, Annika Held, Melina Kurzawe, Ramona Papapostolou, Jade Tessien, Julian Bludau, Andreas Kelch, Sarah Gronefeld, Yannis Schöneberg, Christian Zeitz, Konstantin Zapf, David Prochotta, Maximilian Murphy, Monica M. Sheffer, Moritz Sonnewald, Maria A. Nilsson, Axel Janke

**Affiliations:** Institute for Ecology, Evolution and Diversity, Goethe University, Frankfurt am Main, Germany; Senckenberg Biodiversity and Climate Research Centre, Frankfurt am Main, Germany; LOEWE-Centre for Translational Biodiversity Genomics, Frankfurt am Main, Germany; South African National Biodiversity Institute, National Zoological Garden, Pretoria, South Africa; Zoological Institute and Museum, University of Greifswald, Greifswald, Germany; Senckenberg Research Institute, Department of Marine Zoology, Section Ichthyology, Frankfurt am Main, Germany

**Keywords:** Callionymidae, genome annotation, genome assembly, Hi-C, Nanopore, Syngnathiformes

## Abstract

**Background:** The common dragonet, *Callionymus lyra*, is one of three *Callionymus* species inhabiting the North Sea. All three species show strong sexual dimorphism. The males show strong morphological differentiation, e.g., species-specific colouration and size relations, while the females of different species have few distinguishing characters. *Callionymus* belongs to the ‘benthic associated clade’ of the order Syngnathiformes. The ‘benthic associated clade’ so far is not represented by genome data and serves as an important outgroup to understand the morphological transformation in ‘long-snouted’ syngnatiforms such as seahorses and pipefishes.

**Findings:** Here, we present the chromosome-level genome assembly of *C. lyra*. We applied Oxford Nanopore Technologies’ long-read sequencing, short-read DNBseq, and proximity-ligation-based scaffolding to generate a high-quality genome assembly. The resulting assembly has a contig N50 of 2.2 Mbp, a scaffold N50 of 26.7 Mbp. The total assembly length is 568.7 Mbp, of which over 538 Mbp were scaffolded into 19 chromosome-length scaffolds. The identification of 94.5% of complete BUSCO genes indicates high assembly completeness. Additionally, we sequenced and assembled a multi-tissue transcriptome with a total length of 255.5 Mbp that was used to aid the annotation of the genome assembly. The annotation resulted in 19,849 annotated transcripts and identified a repeat content of 27.66%.

**Conclusions:** The chromosome-level assembly of *C. lyra* provides a high-quality reference genome for future population genomic, phylogenomic, and phylogeographic analyses.

## Data Description

### Background information

Until recently, the family Callionymidae was placed into the order Perciformes, which is often considered a “polyphyletic taxonomic wastebasket for families not placed in other orders” [1]. However, recent phylogenetic analyses suggest a placement of Callionymidae within the order Syngnathiformes, which currently contains ten families with highly derived morphological characters such as the pipefish and seahorses [1]. Syngnathiformes has recently been divided into two clades, a ‘long-snouted clade’ and a ‘benthic associated clade,’ each comprising five families [2]. The ‘long-snouted clade’ (Syngnathidae, Solenostomidae, Aulostomidae, Centriscidae, and Fistulariidae) is currently represented by genomes from the Gulf pipefish (*Syngnathus scovelli*) and the tiger tail seahorse (*Hippocampus comes*) [3,4] and additional draft assemblies of pipefish [5]. A genome of the ‘benthic associated clade’ (Callionymidae, Draconettidae, Dactylopteridae, Mullidae, and Pegasidae) has not been sequenced and analysed yet. Callionymidae comprises 196 species [6], of which the common dragonet*, Callionymus lyra* (Linnaeus, 1758) (Fig. 1), is one of three *Callionymus* species inhabiting the North Sea [7]. All three species also occur in the East Atlantic, and the Mediterranean Sea [6]. They represent essential prey fish for commercially important fish species such as the cod (*Gadus morhua*) [8]. The males of the North Sea dragonet species (*C. lyra, C. maculatus, C. reticulatus*) show strong morphological differentiation in the form of species-specific colouration and size relations. The much less conspicuous females can be distinguished morphologically, with rather high inaccuracy, by the presence or absence of their preopercular, basal spine and by various percentual length ratios. The great resemblance among the different species’ females, together with the fact that all three species can be found in sympatry, suggest the possibility of hybridization among them.

**Figure 1.**
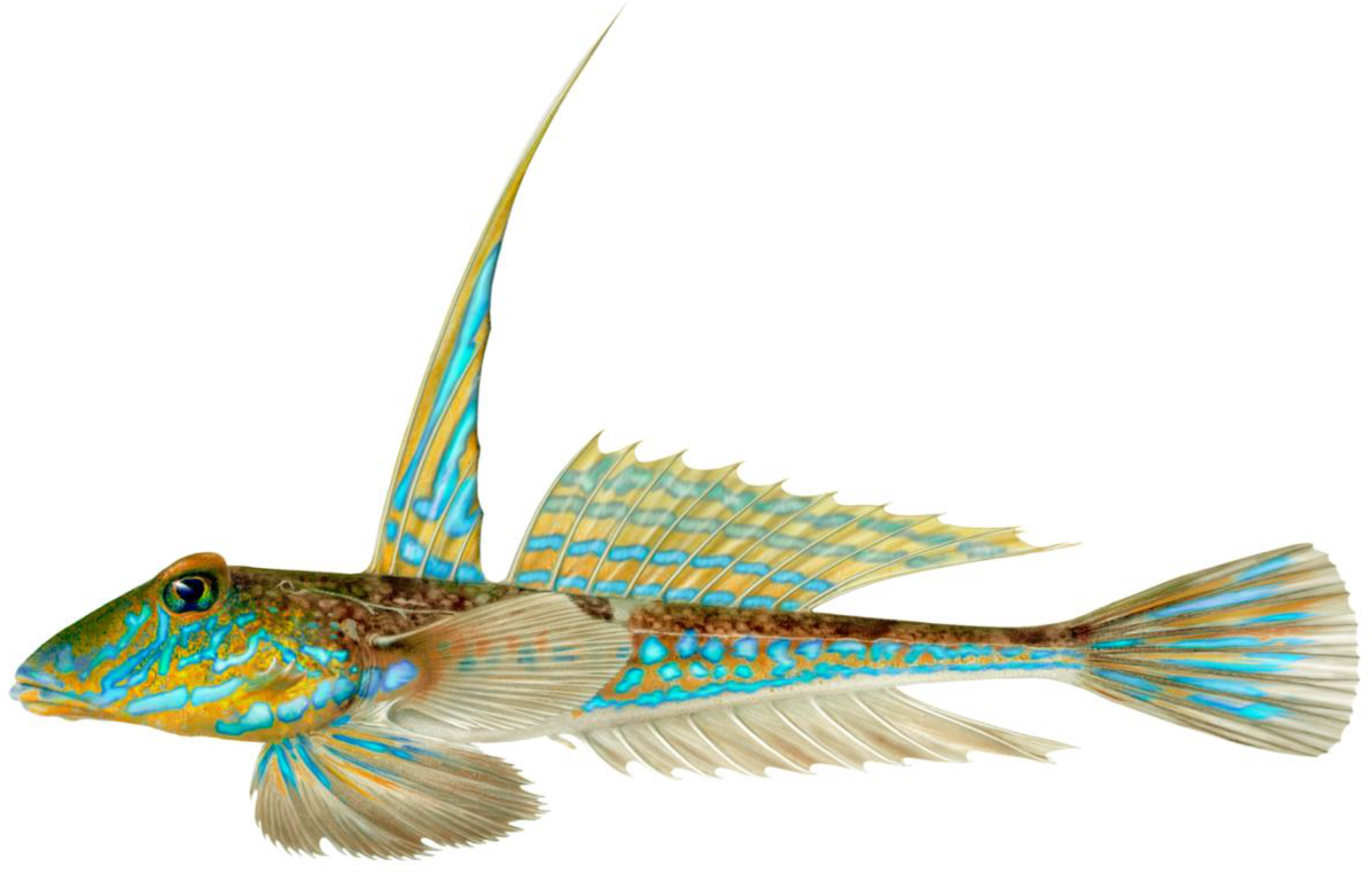
Male *Callionymus lyra*. Artwork by © Karl Jilg/ArtDatabanken

Here, we present the chromosome-level genome of the common dragonet, representing the first genome of the ‘benthic associated’ Syngnathiformes clade as a reference for future population genomic, phylogenomic, and comparative genomic analyses. The chromosome-level genome assembly was generated as part of a six-week university master’s course. For a detailed description and outline of the course, see [9].

### Sampling, DNA extraction, and Sequencing

We sampled two *Callionymus lyra* individuals (one of each sex) during a yearly monitoring expedition to the Dogger Bank in the North Sea (Female: 54°59.189’N 1°37.586’E; Male: 54°48.271’N 1°25.077’E) with the permission of the Maritime Policy Unit of the UK Foreign and Commonwealth Office in 2019. The samples were initially frozen at −20°C on the ship and later stored at −80°C until further processing. The study was conducted in compliance with the ‘Nagoya Protocol on Access to Genetic Resources and the Fair and Equitable Sharing of Benefits Arising from Their Utilization’.

We extracted high molecular weight genomic DNA (hmwDNA) from muscle tissue of the female individual following the protocol by [10]. Quantity and quality of the DNA was evaluated using the Genomic DNA ScreenTape on the Agilent 2200 TapeStation system (Agilent Technologies). Library preparation for long-read sequencing followed the associated protocols for Oxford Nanopore Technologies (ONT, Oxford, UK) Rapid Sequencing Kit (SQK-RAD004). A total of seven sequencing runs were performed using individual flow cells (FLO-MIN106 v.9.41) on a ONT MinION v.Mk1B.

Additionally, we sent tissue samples to BGI Genomics (Shenzhen, China) to generate additional sequencing data. A 100 bp paired-end short-read genomic DNA sequencing library was prepared from the muscle tissue of the female individual. This library was later used for genome assembly polishing. Moreover, a 100 bp paired-end RNAseq library was prepared for pooled RNA isolates derived from kidney, liver, gill, gonad, and brain tissue of the male individual. Both libraries were sequenced on BGI’s DNBseq platform. We received a total of 159,925,221 read pairs (~32 Gbp) of pre-filtered genomic DNA sequencing data and 61,496,990 read-pairs (~12.3 Gbp) of pre-filtered RNAseq data.

Furthermore, we prepared a Hi-C library using the Dovetail™ Hi-C Kit (Dovetail Genomics, Santa Cruz, California, USA) from muscle tissue of the female and sent the library to Novogene Co., Ltd. (Beijing, China) for sequencing on an Illumina NovaSeq 6000. Sequencing yielded a total of 104,668,356 pre-filtered 150 bp paired-end read pairs or 31.4 Gbp of sequencing data. This data was used for proximity-ligation scaffolding of the assembly.

### Genome size estimation

We estimated the genome size for *C. lyra* using both k-mer frequencies and flow cytometry. The k-mer frequency for K=21 was calculated from the short-read DNBseq data and summarized as histograms with jellyfish v.2.2.10 (RRID:SCR_005491) [11]. Plotting the histograms and calculating the genome size and heterozygosity with GenomeScope v.1.0 (RRID:SCR_017014) [12] resulted in a genome size estimate of approximately 562 Mbp. For the genome size estimation using flow cytometry, frozen muscle tissue was finely chopped with a razor blade in 200 μl LeukoSure Lyse Reagent (Beckman Coulter Inc., Fullerton, CA, USA). Large debris was removed by filtering through a 40 μm Nylon cell strainer and an RNAse treatment was performed with a final concentration of 0.3 mg/ml. Simultaneously, we stained the DNA in the nuclei with propidium iodide (PI) at a final concentration of 0.025 mg/ml and incubated the solution for 30 min at room temperature, protected from light exposure. Fluorescence intensities of the nuclei were recorded on the CytoFLEX Flow Cytometer (Beckman Coulter Inc., Fullerton, CA, USA). The domestic cricket (*Acheta domesticus*, C-value: 2.0 pg) was used as a reference to determine the genome size of *C. lyra*. For a more precise estimate we analysed five independent technical replicates resulting in an average C-value of 0.66 pg, which corresponds to a haploid genome size of 645 Mbp.

### Genome assembly and polishing

Nanopore raw signal data (fast5) of the seven sequencing runs were base-called with Guppy v.3.2.4 (ONT) using the high accuracy setting. All individual sequencing runs were examined and compared with NanoComp v.1.0.0 (Supplementary Fig. 1, Supplementary Table 1) [13]. The final dataset, after concatenation of all read-files, was further examined with NanoPlot v.1.0.0 (Supplementary Table 1) [13]. Concatenation of all read-files resulted in a total dataset of 31 Gbp or approximately 55-fold coverage as the basis for the genome assembly.

We assembled the genome of *C. lyra* with wtdbg2 v.2.2 (RRID:SCR_017225) [14] using the default parameters for ONT reads. The resulting assembly was subjected to a three-step polishing approach. First, a single iteration of racon v.1.4.3 (RRID:SCR_017642) [15] corrected for errors typical of the MinION platform: homopolymers and repeat errors. Next, we used one iteration of medaka v.0.11.5 [16] on the racon-polished assembly. According to the developers medaka is most effective after a polishing run with racon. Following polishing with the long-read data, we used three iterations of pilon v.1.23 (RRID:SCR_014731) [17] to correct for random errors and single-base errors with the high-quality short-read data.

### Assembly QC and scaffolding

We calculated assembly continuity statistics using QUAST v.5.0.2 (RRID:SCR_001228) [18] and performed a gene set completeness analysis using BUSCO v.4.0.6 (RRID:SCR_015008) [19] with the provided database for Actinopterygii orthologous genes (actinopterygii_odb10). The final polished assembly had 1,782 contigs and a total length of 569 Mbp, which is marginally larger than the k-mer based estimate of 562 Mbp and 84 Mbp shorter than the flow cytometry estimate. This is expected, because very repetitive regions are usually missing or collapsed in a genome assembly, which could explains the shorter assembly length compared to the flow cytometry size estimate. The assembly shows a high continuity with long contigs of up to 10.7 Mbp and a contig N50 of > 2.2 Mbp (Table 1). The genome assembly completeness analysis identified 95.0 % complete BUSCO genes (93.6% complete, single copy) and only 4.4% missing BUSCOs, which suggests that the assembly contains most of the coding regions of the genome (Fig. 2, Supplementary Table 2).

**Table 1.** Assembly statistics of the long-read based contig assembly (wtdbg2), Hi-C scaffolded assembly (HiRise) and the transcriptome assembly of *Callionymus lyra*.

|  | wtDBG2 | HiRise* | transcriptome |
| --- | --- | --- | --- |
| **No. of contigs** | 1,782 | 1,205 | 246,012 |
| **No. of contigs (> 1 kbp)** | 1,707 | 1,151 | 66,332 |
| **L50** | 70 | 9 | 23,587 |
| **L75** | 176 | 14 | 49,697 |
| **N50** | 2,201,294 bp | 26,698,546 bp | 2,787 bp |
| **N75** | 856,699 bp | 22,283,913 bp | 1,474 bp |
| **Max. contig length** | 10,738,616 bp | 51,234,906 bp | 31,800 bp |
| **Total length** | 569,037,589 bp | 568,707,486 bp | 255,540,591 bp |
| **GC (%)** | 38.97 | 38.97 | 46.84 |
| **No. of gaps** | 0 | 57,348 | 0 |
| **No. of N's per 100 kbp** | 0.0 | 10.08 | 0.0 |

\*final assembly after removing contaminated scaffolds and scaffolds < 200 bp

**Figure 2:**
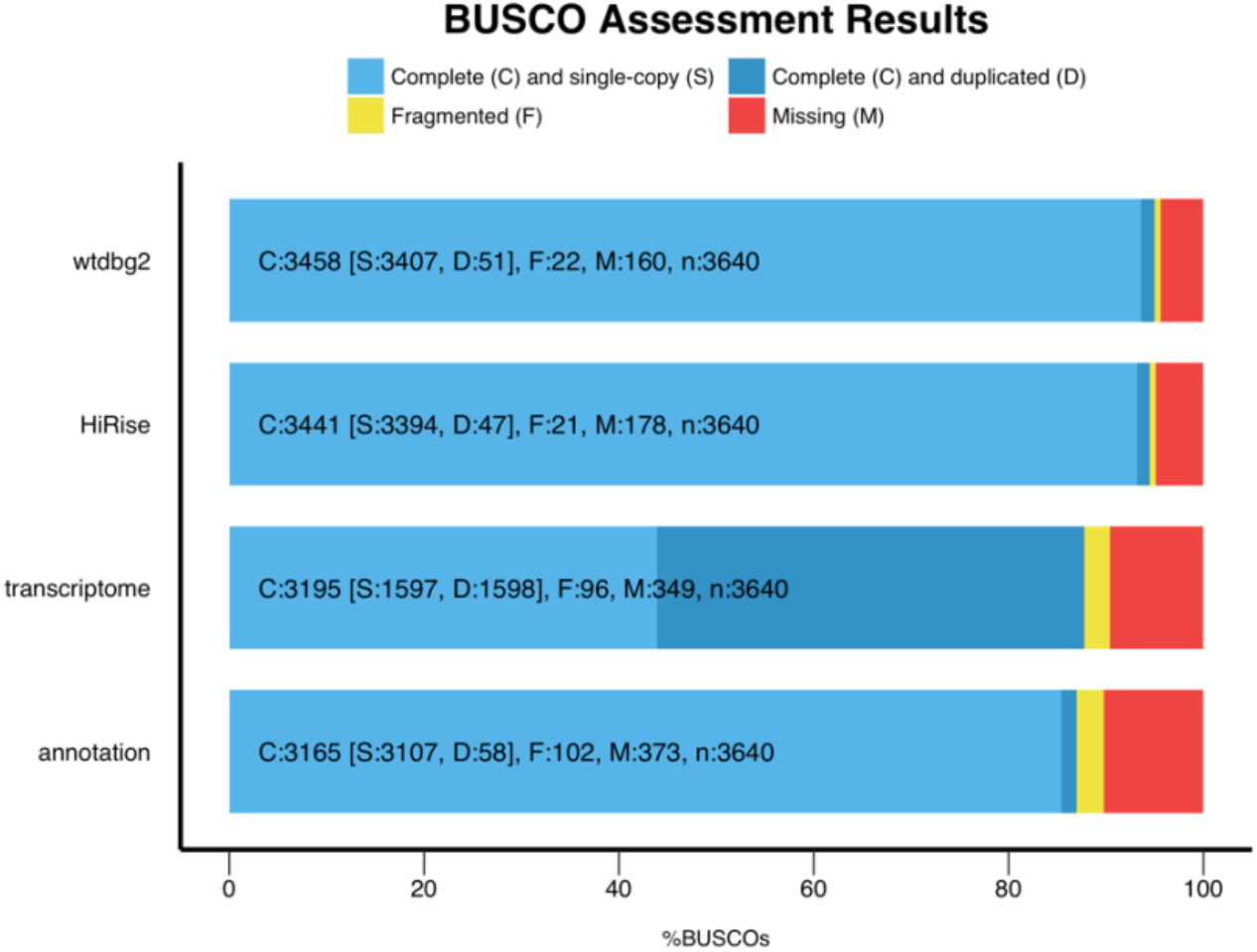
Gene completeness analysis of the long-read based contig assembly (wtdbg2), the Hi-C scaffolded assembly (HiRise), the transcriptome of *Callionymus lyra*, and the annotation. The high percentage of duplicated BUSCOs in the transcriptome is attributed to protein isoforms.

To achieve chromosome-length scaffolds, we used the long-read based assembly and the generated Hi-C data as input for the HiRise scaffolding pipeline [20] as part of the Dovetail Genomics’ scaffolding service. HiRise made 538 joins and 10 breaks resulting in a scaffolded assembly with a total of 1,254 scaffolds and a scaffold N50 of 26.7 Mbp. Over 94.5% (538 Mbp) of the total assembly length was scaffolded into 19 chromosome-length scaffolds (Fig. 3A). The number of chromosome-length scaffolds is consistent with the haploid number of chromosomes derived from karyotypes of females of two Callionymidae species (*C. beniteguri* and *Repomucenus ornatipinnis*) [21]. Therefore, the number of chromosomes appears to be relatively conserved within *Callionymidae* and it is likely that *C. lyra* follows the same chromosomal sex determination system as *C. beniteguri* and *R. ornatipinnis* (♀: X_1_X_2_-X_1_X_2_ (2n = 38); ♂: X_1_X_2_-Y (2n = 37)) [21].

**Figure 3:**
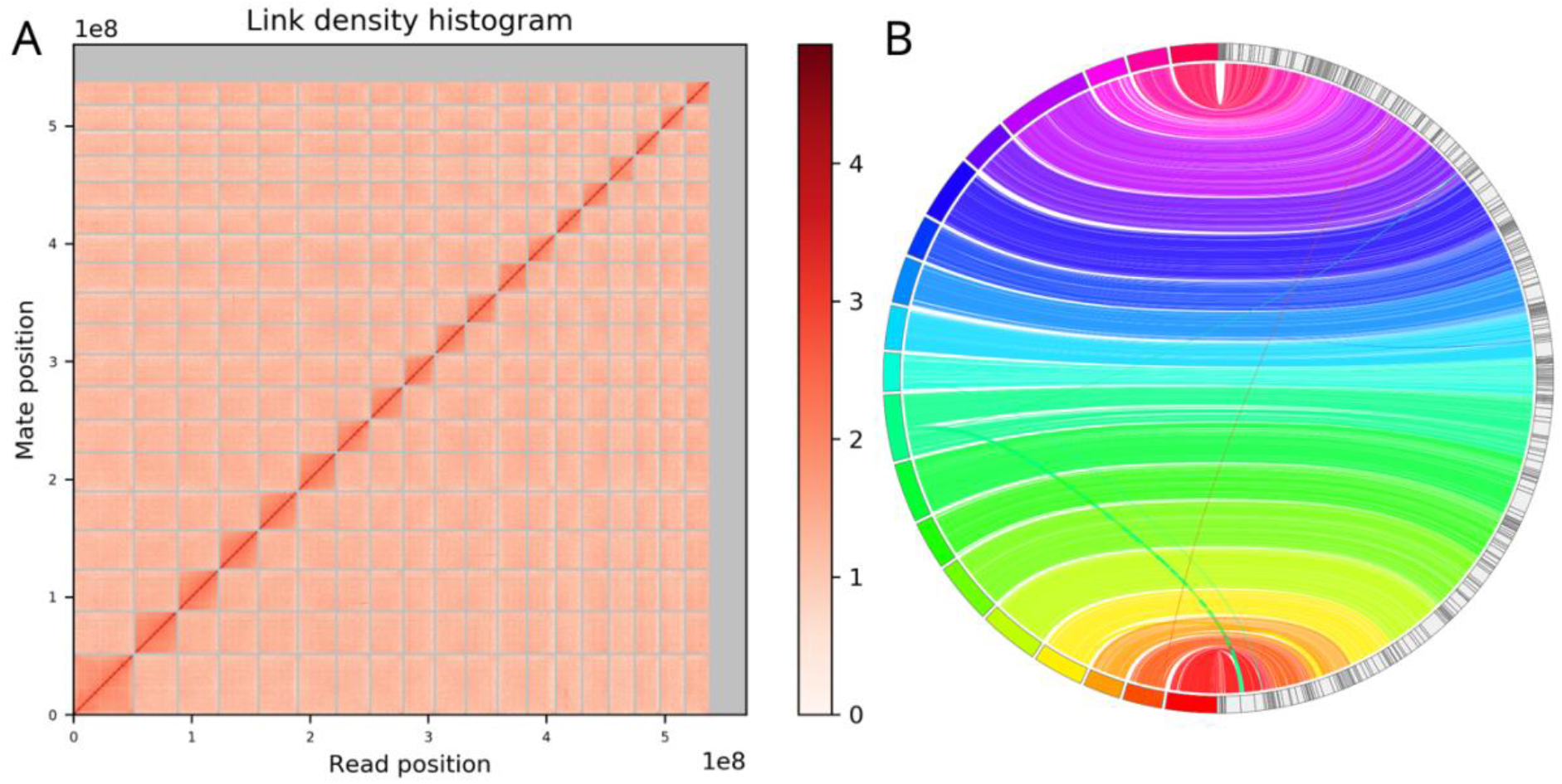
(A) Hi-C contact map of the 19 chromosome-length scaffolds, and additional unplaced scaffolds. (B) Whole genome synteny between the polished contig assembly from wtdbg2 (on the right) and the final Hi-C scaffolded chromosome-level assembly (on the left). Crossing lines indicate assembly artifacts corrected during scaffolding.

For a final assembly quality control, we mapped the raw nanopore reads with minimap2 v.2.17-r941 [22] and the DNBSeq data with bwa-mem v.0.7.17-r1194-dirty (RRID:SCR_010910) [23] onto the final assembly with a high mapping rate of 94.8% and 98.62%, respectively. We further checked the assembly for contamination with BlobTools v.1.1.1 (RRID:SCR_017618) [24]. This analysis identified minor contamination from Proteobacteria (26 short contigs, in total 0.25 Mbp) and Uroviricota (2 contigs, in total 0.12 Mbp) (Supplementary Fig. 2). No contamination was found in the 19 chromosome-length scaffolds. Subsequently, we removed all contaminations and contigs with a length of < 200 bp from the final assembly (for final statistics see Table 1). In addition, we screened for mitochondrial sequence contamination with BLASTN v.2.9.0+ (RRID:SCR_001598) [25] using the available mitochondrial genome sequence of *C. lyra* (Accession No.: MN122938.1) as a reference. A single sequence of mitochondrial origin (169 bp) was identified on one scaffold. This partial mitochondrial sequence could either be an assembly artifact or nuclear mitochondrial DNA (numt).

A synteny plot comparing the polished wtdbg2 contig assembly with the final chromosome-level assembly, generated with JupiterPlot v.1.0 [26], found overall strong agreements with only few differences (Fig. 3B). These likely constitute assembly errors in the contig assembly that were fixed by HiRise during scaffolding. A BUSCO analysis of the final assembly found slightly less complete BUSCO genes compared to the wdtbg2 contig assembly (94.5% vs. 95.0%) (Fig. 2, Supplementary Table 2).

### Transcriptome assembly and Quality

In addition to the genome, we assembled the transcriptome of *C. lyra* for subsequent use in the genome annotation using Trinity v.2.9.0 (RRID:SCR_013048) [27,28] based on the 12.3 Gbp multi-tissue RNAseq data. The resulting transcriptome assembly has a total length of 255.5 Mbp (Table 1). BUSCO analysis suggests a high transcriptome completeness with 87.8% of orthologous genes found in the transcriptome assembly (Fig. 2, Supplementary Table 2).

### Genome annotation

#### Repeat annotation

In order to annotate repeats in the assembly, we created a custom *de novo* repeat library using RepeatModeler v.1.0.11 (RRID:SCR_015027) [29] and combined this library with the *Actinopterygii* repeat database from RepBase. Repeats in the genome were then annotated using RepeatMasker open-4.0.7 (RRID:SCR_012954) [30]. Our analyses identified 27.66% of repeats in the genome, of which the majority consisted of DNA transposons (6.10%), LINE’s (5.32%) and simple repeats (3.47%). Additionally, 10.69% of unclassified repeats were identified (Table 2).

**Table 2.**
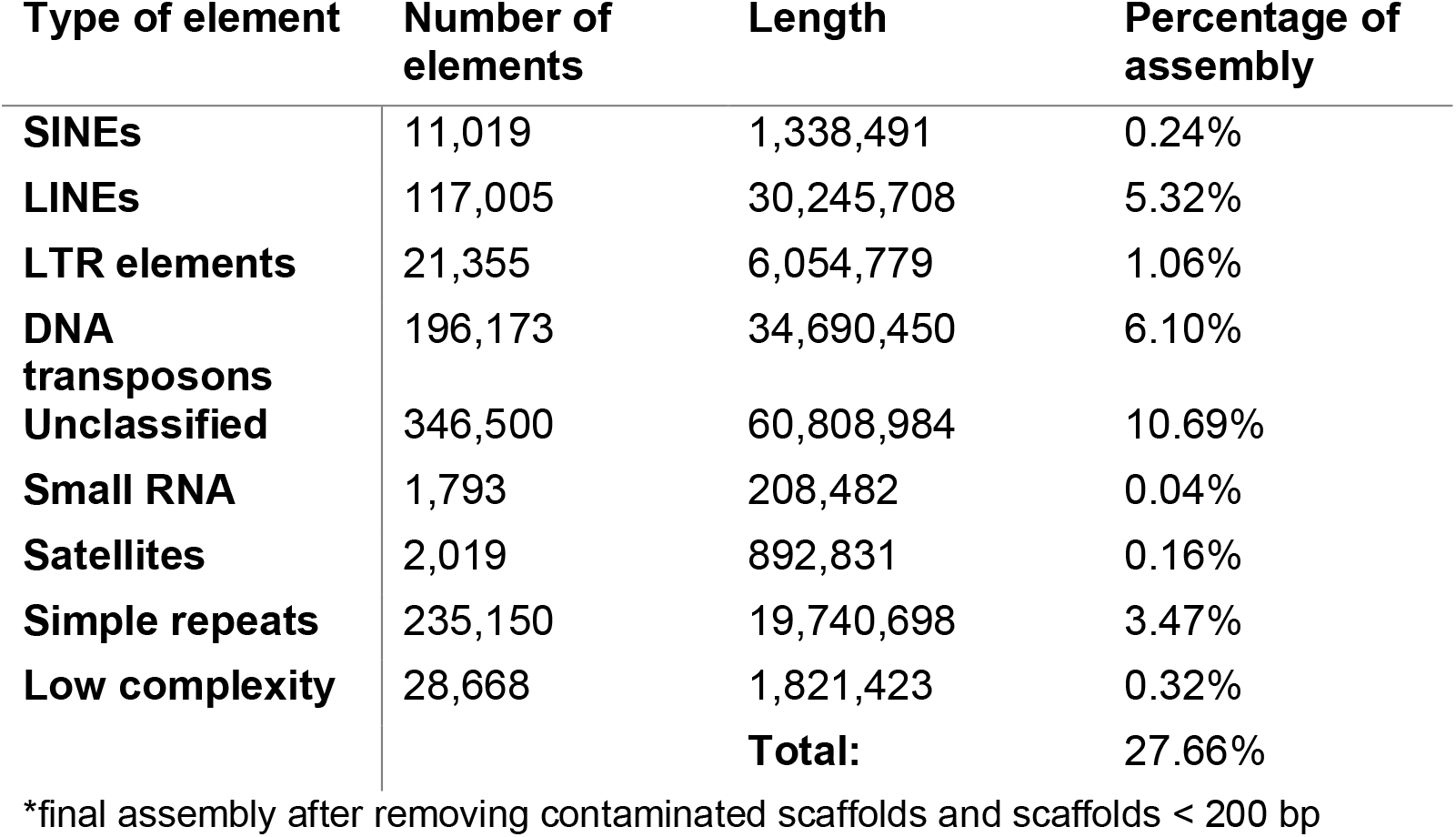
Repeat content of the Hi-C scaffolded assembly*.

#### Gene Annotation

Prior to annotating genes, interspersed repeats in the genome were hard-masked and simple repeats soft-masked to increase the accuracy and efficiency of locating genes. Gene annotation was performed using MAKER2 v.2.31.10 (RRID:SCR_005309) [31]. First, evidence-based annotation was conducted based on the *de novo* assembled transcriptome and previously published proteins from SwissProt [32], and the tiger tail seahorse (*Hippocampus comes*) [3]. Next, genes were *ab initio* predicted with SNAP v.2006-07-28 (RRID:SCR_002127) [33] and Augustus v.3.3 (RRID:SCR_008417) [34]. The final gene annotation resulted in 19,849 transcripts, of which 96% had an AED score of ≤ 0.5 (Supplementary Fig. 3), indicating a high quality of the annotated gene models [35].

## Conclusion

Here we report the first genome assembly of the ‘benthic associated’ Syngnathiformes clade, the sister group to the ‘long-snouted clade’ (e.g., seahorses and pipefish). The annotated genome of *Callionymus lyra*, with its high continuity (chromosome-level), provides an essential reference to study speciation and potential hybridization in Callionymidae and is an important resource for phylogenomic analyses among syngnathiform fish.

## Availability of supporting data

All raw data generated in this study including Nanopore long-reads, DNBSeq short-reads, Hi-C reads, and RNASeq data, and the chromosome-level assembly are accessible at GenBank under BioProject PRJNA634838.

## Acknowledgements

We thank Damian Baranski from the TBG laboratory centre for laboratory support. The present study is a result of the Centre for Translational Biodiversity Genomics (LOEWE-TBG) and was supported through the programme “LOEWE – Landes-Offensive zur Entwicklung Wissenschaftlich-ökonomischer Exzellenz” of Hesse’s Ministry of Higher Education, Research, and the Arts.

## Author Contributions

SW, SP, MAN, MS, and AJ designed the study. SW, MN, AH, MK, RP, JT, JB, AK, SG, YS, CZ, KZ, DP, MM, and MMS performed laboratory procedures and sequencing. SP, JDR, RC, MW, MAN, MN, AH, MK, RP, JT, JB, AK, SG, YS, CZ, KZ, DP, MM, and SW conducted bioinformatic processing and analyses. All authors contributed to writing this manuscript.

## List of abbreviations

BLASTN: Basic Local Alignment Search Tool (for nucleotides)
bp: base pairs
BUSCO: Benchmarking Universal Single-Copy Orthologs
DNBSeq: DNA NanoBall sequencing
Gbp: Gigabase pairs
hmwDNA: high molecular weight DNA
kbp: kilobase pairs
Mbp: megabase pairs
numt: nuclear mitochondrial DNA
ONT: Oxford Nanopore Technologies
pg: picogram
PI: propidium iodide
RNAseq: RNA sequencing

## Conflict of Interest

The authors declare that they have no competing interests.

## Consent for publication

Not Applicable

## Notes

### Competing Interest Statement

The authors have declared no competing interest.

## References

1. Betancur-R R, Wiley EO, Arratia G, Acero A, Bailly N, Miya M, et al. Phylogenetic classification of bony fishes. BMC Evol Biol. 2017;17:162.

2. Longo SJ, Faircloth BC, Meyer A, Westneat MW, Alfaro ME, Wainwright PC. Phylogenomic analysis of a rapid radiation of misfit fishes (Syngnathiformes) using ultraconserved elements. Mol Phylogenet Evol. 2017;113:33–48.

3. Lin Q, Fan S, Zhang Y, Xu M, Zhang H, Yang Y, et al. The seahorse genome and the evolution of its specialized morphology. Nature. Nature Publishing Group; 2016;540:395–9.

4. Small CM, Bassham S, Catchen J, Amores A, Fuiten AM, Brown RS, et al. The genome of the Gulf pipefish enables understanding of evolutionary innovations. Genome Biol. 2016;17:258.

5. Roth O, Solbakken MH, Tørresen OK, Bayer T, Matschiner M, Baalsrud HT, et al. Evolution of male pregnancy associated with remodeling of canonical vertebrate immunity in seahorses and pipefishes. Proc Natl Acad Sci. National Academy of Sciences; 2020;117:9431–9.

6. Froese R, Pauly D. FishBase [Internet]. World Wide Web Electron. Publ. Version 122019. 2019. Available from: www.fishbase.org

7. Fricke R. Annotated checklist of the dragonet families Callionymidae and Draconettidae (Teleostei: Callionymoidei), with comments on callionymid fish classification. Stuttg Beitr Naturk Ser Biol. 2002;645:1–103.

8. Armstrong MJ. The predator-prey relationships of Irish Sea poor-cod (*Trisopterus minutus* L.), pouting (*Trisopterus luscus* L.) and cod (*Gadus morhua* L.). ICES J Mar Sci. Oxford Academic; 1982;40:135–52.

9. Prost S, Winter S, De Raad J, Coimbra RTF, Wolf M, Nilsson MA, et al. Education in the genomics era: Generating high-quality genome assemblies in university courses. GigaScience. 2020;9.

10. Mayjonade B, Gouzy J, Donnadieu C, Pouilly N, Marande W, Callot C, et al. Extraction of high-molecular-weight genomic DNA for long-read sequencing of single molecules. BioTechniques. 2016;61:203–5.

11. Marçais G, Kingsford C. A fast, lock-free approach for efficient parallel counting of occurrences of k-mers. Bioinformatics. 2011;27:764–70.

12. Vurture GW, Sedlazeck FJ, Nattestad M, Underwood CJ, Fang H, Gurtowski J, et al. GenomeScope: fast reference-free genome profiling from short reads. Bioinformatics. 2017;33:2202–4.

13. De Coster W, D’Hert S, Schultz DT, Cruts M, Van Broeckhoven C. NanoPack: visualizing and processing long-read sequencing data. Bioinformatics. 2018;34:2666–9.

14. Ruan J, Li H. Fast and accurate long-read assembly with wtdbg2. Nat Methods. 2020;17:155–8.

15. Vaser R, Sović I, Nagarajan N, Šikić M. Fast and accurate de novo genome assembly from long uncorrected reads. Genome Res. 2017;27:737–46.

16. medaka [Internet]. Oxford Nanopore Technologies; 2020 [cited 2020 Jul 23]. Available from: https://github.com/nanoporetech/medaka

17. Walker BJ, Abeel T, Shea T, Priest M, Abouelliel A, Sakthikumar S, et al. Pilon: An Integrated Tool for Comprehensive Microbial Variant Detection and Genome Assembly Improvement. PLOS ONE. 2014;9:e112963.

18. Gurevich A, Saveliev V, Vyahhi N, Tesler G. QUAST: quality assessment tool for genome assemblies. Bioinformatics. 2013;29:1072–5.

19. Seppey M, Manni M, Zdobnov EM. BUSCO: Assessing Genome Assembly and Annotation Completeness. Methods Mol Biol Clifton NJ. 2019;1962:227–45.

20. Putnam NH, O’Connell BL, Stites JC, Rice BJ, Blanchette M, Calef R, et al. Chromosome-scale shotgun assembly using an in vitro method for long-range linkage. Genome Res. 2016;26:342–50.

21. Murofushi M, Nishikawa S, Yosida TH. Cytogenetical Studies on Fishes. V. Proc Jpn Acad Ser B. 1983;59:58–61.

22. Li H. Minimap2: pairwise alignment for nucleotide sequences. Bioinformatics. 2018;34:3094–100.

23. Li H. Aligning sequence reads, clone sequences and assembly contigs with BWA-MEM. ArXiv Prepr ArXiv13033997. 2013;

24. Laetsch DR, Blaxter ML. BlobTools: Interrogation of genome assemblies. F1000Research. 2017;6:1287.

25. Zhang Z, Schwartz S, Wagner L, Miller W. A Greedy Algorithm for Aligning DNA Sequences. J Comput Biol. Mary Ann Liebert, Inc., publishers; 2000;7:203–14.

26. Chu J. Jupiter Plot: A Circos-based tool to visualize genome assembly consistency. 2018; Available from: http://doi.org/10.5281/zenodo.1241235

27. Grabherr MG, Haas BJ, Yassour M, Levin JZ, Thompson DA, Amit I, et al. Full-length transcriptome assembly from RNA-Seq data without a reference genome. Nat Biotechnol. 2011;29:644–52.

28. Haas BJ, Papanicolaou A, Yassour M, Grabherr M, Blood PD, Bowden J, et al. De novo transcript sequence reconstruction from RNA-seq using the Trinity platform for reference generation and analysis. Nat Protoc. 2013;8:1494–512.

29. RepeatModeler [Internet]. [cited 2020 Jul 23]. Available from: http://www.repeatmasker.org/RepeatModeler/

30. RepeatMasker [Internet]. [cited 2020 Jul 23]. Available from: http://www.repeatmasker.org/RMDownload.html

31. Holt C, Yandell M. MAKER2: an annotation pipeline and genome-database management tool for second-generation genome projects. BMC Bioinformatics. 2011;12:491.

32. The UniProt Consortium. UniProt: a worldwide hub of protein knowledge. Nucleic Acids Res. Oxford Academic; 2019;47:D506–15.

33. Korf I. Gene finding in novel genomes. BMC Bioinformatics. 2004;5:59.

34. Stanke M, Diekhans M, Baertsch R, Haussler D. Using native and syntenically mapped cDNA alignments to improve de novo gene finding. Bioinformatics. 2008;24:637–44.

35. Yandell M, Ence D. A beginner’s guide to eukaryotic genome annotation. Nat Rev Genet. Nature Publishing Group; 2012;13:329–42.

