## Supplementary material for "Chromosome-level genome assembly of a benthic associated Syngnathiformes species: the common dragonet, *Callionymus lyra*"

### Title

### Affiliations

Supplementary Figures:

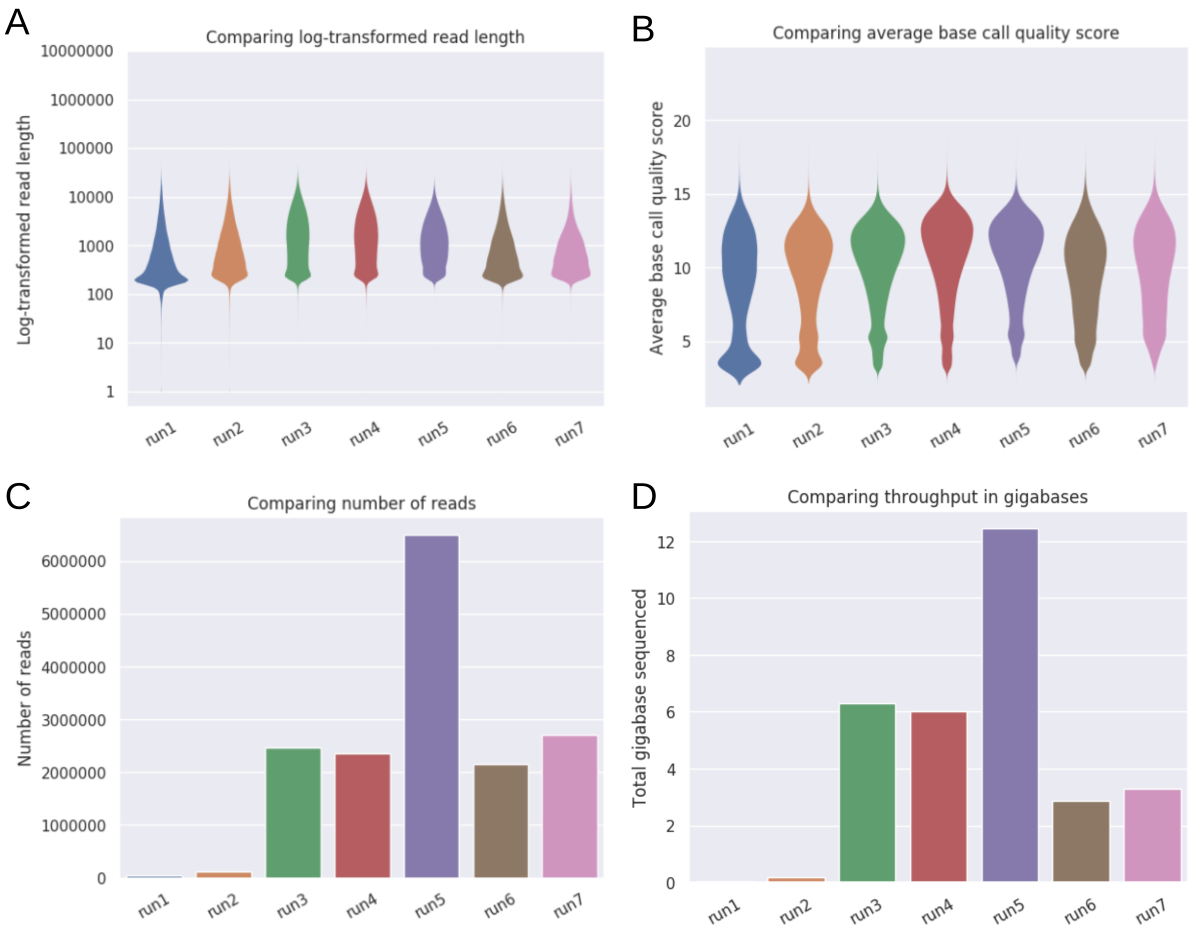


**Supplementary Figure 1**.
Comparison of the data output and read lengths between all seven MinION sequencing runs. All libraries were prepared with the ONT Rapid sequencing kit (SQK-RAD004). A) log-transformed read lengths per run, B) Read quality scores of the seven different runs, C) numbers of reads, and D) the total amount of sequencing data generated per run.

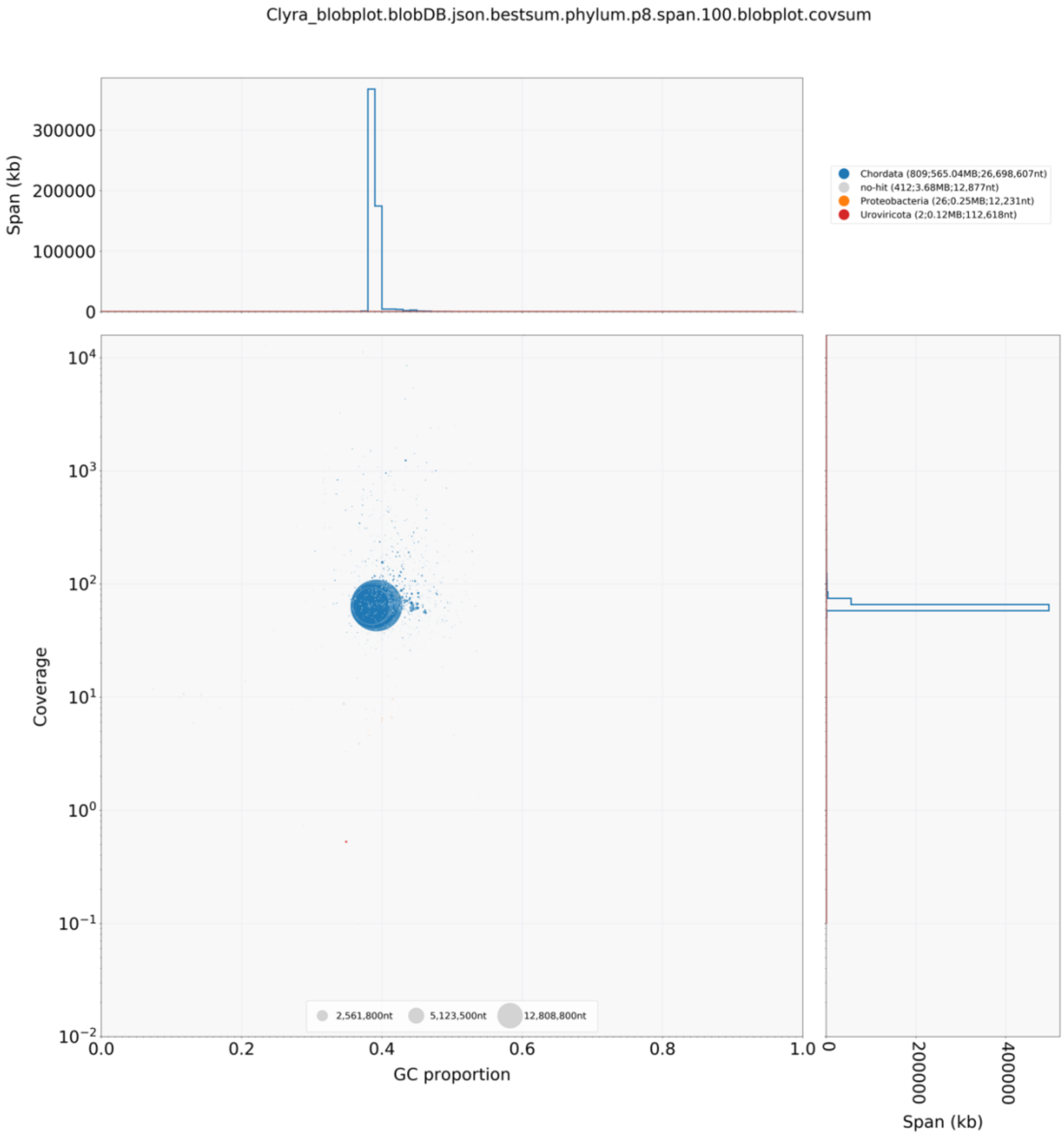


**Supplementary Figure 2.**

Blobtools plot showing the taxonomic assignments (blue colour for Chordata, gray for “no hits”, orange for Proteobacteria, and red for Uroviricota) of the different scaffolds, and scaffold-wide coverage and GC contents. The scaffolds were blasted against the NCBI nucleotide database.

Scaffolds with assignments to Proteobacteria or Uroviricota were removed from the final assembly.

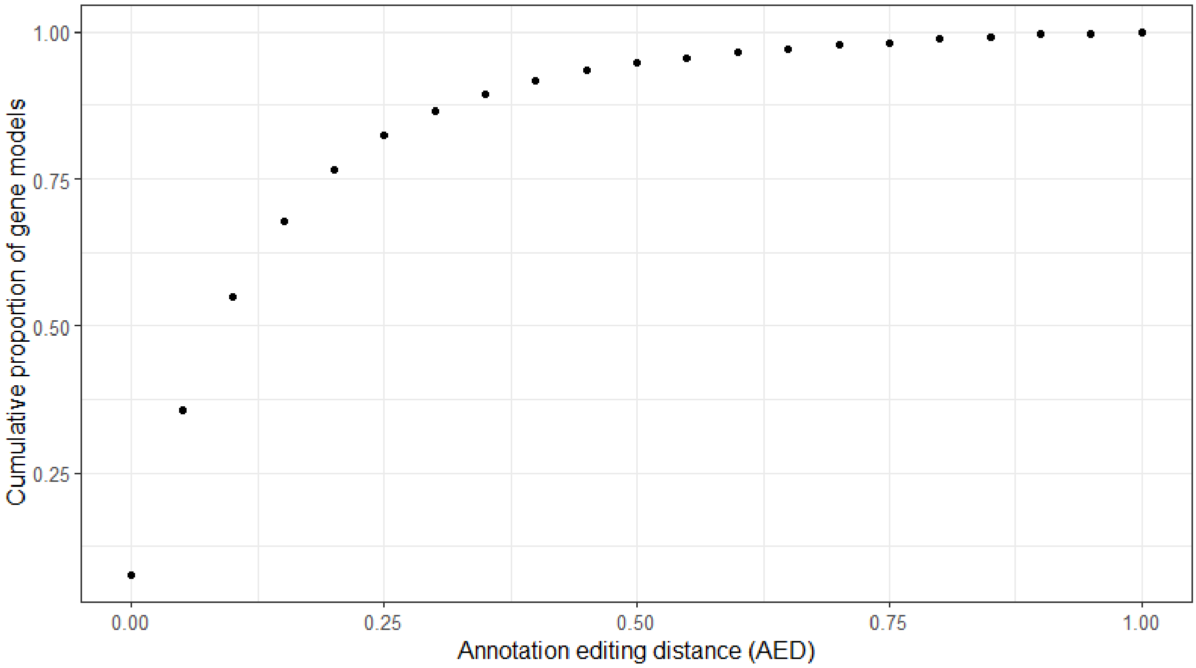


**Supplementary Figure 3.**
Distribution of Annotation Edit Distance (AED) scores. About 96% of all gene models show AED scores of ≤ 0.5 indicating a high quality of the gene models.

Supplementary Tables:

**Supplementary Table 1. Read output and quality of the seven different MinION sequencing runs and the final concatenated dataset.**

| | Run 1 | Run 2 | Run 3 | Run 4 | Run 5 | Run 6 | Run 7 | total |
|---|---|---|---|---|---|---|---|---|
| **Mean read length** | 1,153 | 1,528 | 2,562 | 2,542 | 1,913 | 1,334 | 1,211 | 1,904 |
| **Mean read quality:** | 8.7 | 9.3 | 10.1 | 10.6 | 10.6 | 9.4 | 9.9 | 10.2 |
| **Number of reads:** | 49,659 | 114,845 | 2,465,768 | 2,360,176 | 6,506,852 | 2,149,172 | 2,714,852 | 16,361,324 |
| **Read length N50:** | 3,425 | 3,628 | 5,469 | 5,257 | 3,485 | 2,880 | 2,179 | 3,931 |
| **Total bases:** | 57,260,945 | 175,534,110 | 6,316,560,588 | 5,998,757,646 | 12,447,462,241 | 2,865,915,086 | 3,287,474,510 | 31,148,965,126 |

**Supplementary Table 2.**
BUSCO results of the long-read based contig assembly (wtdbg2), Hi-C scaffolded assembly (HiRise), the transcriptome, and the annotation of the *Callionymus lyra* assembly.

| | **wtdbg2** | **HiRise*** | **transcriptome** | **annotation** |
|---|---|---|---|---|
| Complete BUSCOs | 3458 (95.0%) | 3441 (94.5%) | 3195 (87.8%) | 3165 (87.0%) |
| Complete and single-copy BUSCOs | 3407 (93.6%) | 3394 (93.2%) | 1597 (43.9%) | 3107 (85.4%) |
| Complete and duplicated BUSCOs | 51 (1.4%) | 47 (1.3%) | 1598 (43.9%) | 58 (1.6%) |
| Fragmented BUSCOs | 22 (0.6%) | 21 (0.6%) | 96 (2.6%) | 102 (2.8%) |
| Missing BUSCOs | 160 (4.4%) | 178 (4.9%) | 349 (9.6%) | 373 (10.2%) |
| Total BUSCO groups searched | 3640 | 3640 | 3640 | 3640 |
| *final assembly after removing contaminated scaffolds and scaffolds < 200 bp | | | | |
